# Forest canopy resists plant invasions: a case study of *Chromolaena odorata* in sub-tropical Sal (*Shorea robusta*) forests of Nepal

**DOI:** 10.1101/747287

**Authors:** LN Sharma, B Adhikari, MF Watson, B Karna, E Paudel, BB Shrestha, DP Rijal

**Affiliations:** ForestAction, Nepal, Bagdol Ringroad, Lalitpur, Nepal; Royal Botanic Garden Edinburgh (RBGE), 20a Inverleith Row, Scotland, UK, EH3 5LR; Nepal Academy of Science and Technology, Khumaltar, Lalitpur, Nepal; Central Department of Botany, Tribhuvan University, Kirtipur, Kathmandu, Nepal; Department of Arctic and Marine Biology, UiT: The Arctic University of Norway, Tromsø, Norway

**Keywords:** Biotic resistance, canopy cover, disturbance, forest management, invasive alien species

## Abstract

Invasive Alien Species cause tremendous ecological and economic damage in agriculture, forestry, aquatic ecosystems, and pastoral resources. They are one of the major threats to biodiversity conservation. Understanding the spatial pattern of invasive species and disentangling the biophysical drivers of invasion at forest stand level is essential for managing invasive species in forest ecosystems and the wider landscape. However, forest-level and species-specific information on invasive species abundance and area of extent is largely lacking. In this context, we analysed the cover of one of the world’s worst invasive plant species *Chromolaena odorata* in Sal (*Shorea robusta* Gaertn.) forest in central Nepal. Vegetation was sampled in four community-managed forests using 0.01 ha square quadrats, covering forest edge to the interior. *Chromolaena* cover, floral richness, tree density, forest canopy cover, shrub cover, and tree basal area were measured in each plot. We also estimated the level of disturbance in plots, and calculated distance from the plot to the nearest road. We also explored forest and invasive species management practices in community forests.

*Chromolaena* cover was found to be negatively correlated with forest canopy cover, distance to the nearest road, angle of slope and shrub cover. Canopy cover had the greatest effect on the *Chromolaena* cover. *Chromolaena* cover did not show any pattern along native species richness gradients. In conclusion, forest canopy cover is the overriding biotic covariate affecting *Chromolaena* cover in Sal forests. The practical application of our results in managing *Chromolaena* in forest ecosystems is discussed.

## Introduction

Invasive Alien Species (IAS) are an important component - both driver and passenger - of global environmental change (Vitousek et al. 1997). They are among the major threats to biodiversity and have already caused tremendous economic loss and ecological damage across ecosystems and geographical scales (Millenium Ecosystem Assessment 2005, Pyšek and Richardson 2010, Pyšek et al. 2012). These threats are ever growing with expanding transportation networks and increased mobility of people and commodities (Simberloff et al. 2013, Sardain et al. 2019). IAS compete with native biota, alter and homogenize forest composition, change ecosystem functions, compromise ecosystem services and reduce native species diversity (Bellingham et al. 2018). They also degrade habitat quality for wildlife (Murphy et al. 2013), and potentially impact all types of ecosystems and individual species. Nevertheless, impacts are contingent on the trait of the invading species and ecosystem types exposed to the invasion (Martin et al. 2009, Pyšek et al. 2012, Liebhold et al. 2017).

IAS distribution at larger geographical scales is the result of interplay between ecological and social variables, including transportation networks, national gross domestic production, and population density (Liu et al. 2005, Hulme 2009, Niemiec et al. 2018, Sardain et al. 2019). Road networks and mobility of people not only transport propagules from one place to another, but also create locally disturbed areas which are suitable for propagules to establish (González-Moreno et al. 2014). These factors are fundamental to the early stage of invasion, however, further augmentation of invasive species is determined by local environmental factors including habitat disturbance (Stohlgren et al. 2006).

IAS distribution and abundance may vary substantially at different spatial scales (Foxcroft et al. 2009). Distribution patterns generated from coarse scale spatial data, and models based on climatic suitability may not depict the local scale distribution and abundance of invasive species. Some areas of forests, for example canopy gaps and forest margins, provide more conducive environments than forest interiors for invasion success (Arellano-Cataldo and Smith-Ramírez 2016, Driscoll et al. 2016). Understanding the drivers of local scale patterns of IAS distribution and abundance, therefore, is crucial for the management of invasive species at the site level (Foxcroft et al. 2009).

At local scale, invasion success depends on the interactions between species invasiveness, habitat invasibility and propagule pressure (Lozon and MacIsaac 1997, Dyderski and Jagodziński 2018). Anthropogenic disturbances are important drivers of plant invasions as they facilitate invasion by altering resource availability and competition (Lozon and MacIsaac 1997). Disturbance is often associated with road networks and human population density. Roads, trails, rivers, and streams work as dispersal conduits helping to disperse propagules through the forests. Crossing a dispersal barrier and arriving in a new locality does not necessarily guarantee invasion success. Newly arrived propagules of IAS need to pass through edaphic, microclimatic and biological filters for successful invasion (Mitchell et al. 2006, Theoharides 2007, Gallien et al. 2015).

Resident ecological communities naturally tend to resist the establishment and spread of incoming species, a phenomenon termed ‘biological resistance’ (Levine et al. 2004, Nunez-Mir et al. 2017). It has been hypothesized that species-rich communities have a lower vulnerability to invasion at the local scale (Levine et al. 2004), however, the diversity resistance hypothesis is not always supported by empirical studies (Byun and Lee 2018, Smith and Côté 2019). Rather there are also instances of congruence of higher diversity and higher invasion particularly at larger spatial scales (Stohlgren et al. 2006).

The main mechanism behind the biotic resistance is competition (Nunez-Mir et al. 2017). Competition for key resources, for example light, water and nutrients, and space between incoming species and recipient community may be the main mode of the interactions. The attributes of resident communities that curtail the key resources required for incoming species may vary across resident communities and incoming species. Nevertheless, higher species richness of a native community does not necessarily make these communities more competitive and invasion resistant (Levine 2000, Fridley et al. 2007). Therefore, besides species richness, other attributes of these communities, for example density, crowding and biomass, may make communities more competitive and resistant to invasion (Kennedy et al. 2002, Luo et al. 2018, MacLaren et al. 2019). In forest stands, native species richness, tree density, canopy cover and shrub/sapling layer are important community attributes of invasion resistance (Gómez et al. 2019). These attributes indeed determine the availability of empty niches for successful invasions. Forest stand attributes maybe relatively more important than other local factors for invasion success on forest floor by limiting the amount of light reaching the surface of the ground (Charbonneau and Fahrig 2004, Fajardo and Gundale 2018, Bustamante et al. 2019). Shrub/saplings and ground vegetation layers potentially reinforce the impacts of canopy cover by preventing intercepted light falling on the ground. Nevertheless, the impact of canopy may also be dependent on the nature of invading species, shade tolerant invasive species being favored in dense forest stands (Martin et al. 2009).

Distribution and abundance of invasive species at forest stand-level and their ecological correlates are the basic knowledge necessary for the management of invasive species at forest stand level. How attributes of forest stand influence the abundance of IAS is key to managing forest ecosystems sustainably. However, such information and fine scale ecological correlates are rarely available for specific forest types and species. Sal (*Shorea robusta* Gaertn.) forest, a major dominant forest type in the tropical and subtropical parts of the Indian subcontinent, has been heavily impacted by invasion by *Chromolaena odorata* (L.) R.M.King & H.Rob. (hereafter referred to as *Chromolaena*). It has been reported that the abundance of this species shows a positive correlation to forest disturbances and light intensity (Joshi 2006). Nevertheless, this inference was drawn from study conducted on large-sized plot, therefore this study sets out to analyse how forest stand attributes, including canopy cover, influences *Chromolaena* coverage in Sal forest using small-sized multiple plots across a canopy cover gradient. This study seeks to establish how abundance of one of the world’s worst IAS is affected by forest stand attributes (canopy cover, tree density, basal area and shrub cover) and plot level disturbances. We also use this research to test the hypothesis that plots richer in native species are more resist to invasion.

## Methodology

### Study area

We conducted this study in four community-managed forests (CFs) of central Nepal; two in each of Makawanpur and Nawalparasi districts. All the sampled forests were similar in terms of geography, climate, vegetation and management regime; however the forests in Nawalparasi were more fragmented than in Makawanpur (Figure 1). Among them, Pashupati CF was the smallest 167 hectare (ha) and Sunachuri was the largest 260 ha, Janakalyan and Ghumauri are 182 and 208 ha respectively. CFs are forest categories that are managed by local users formed into legally recognised organizations. Nepal has exemplary success in the sustainable management of forest commons through its CF program, with over 20,000 CF User Groups (CFUGs) have been formed and registered.

**Figure 1:**
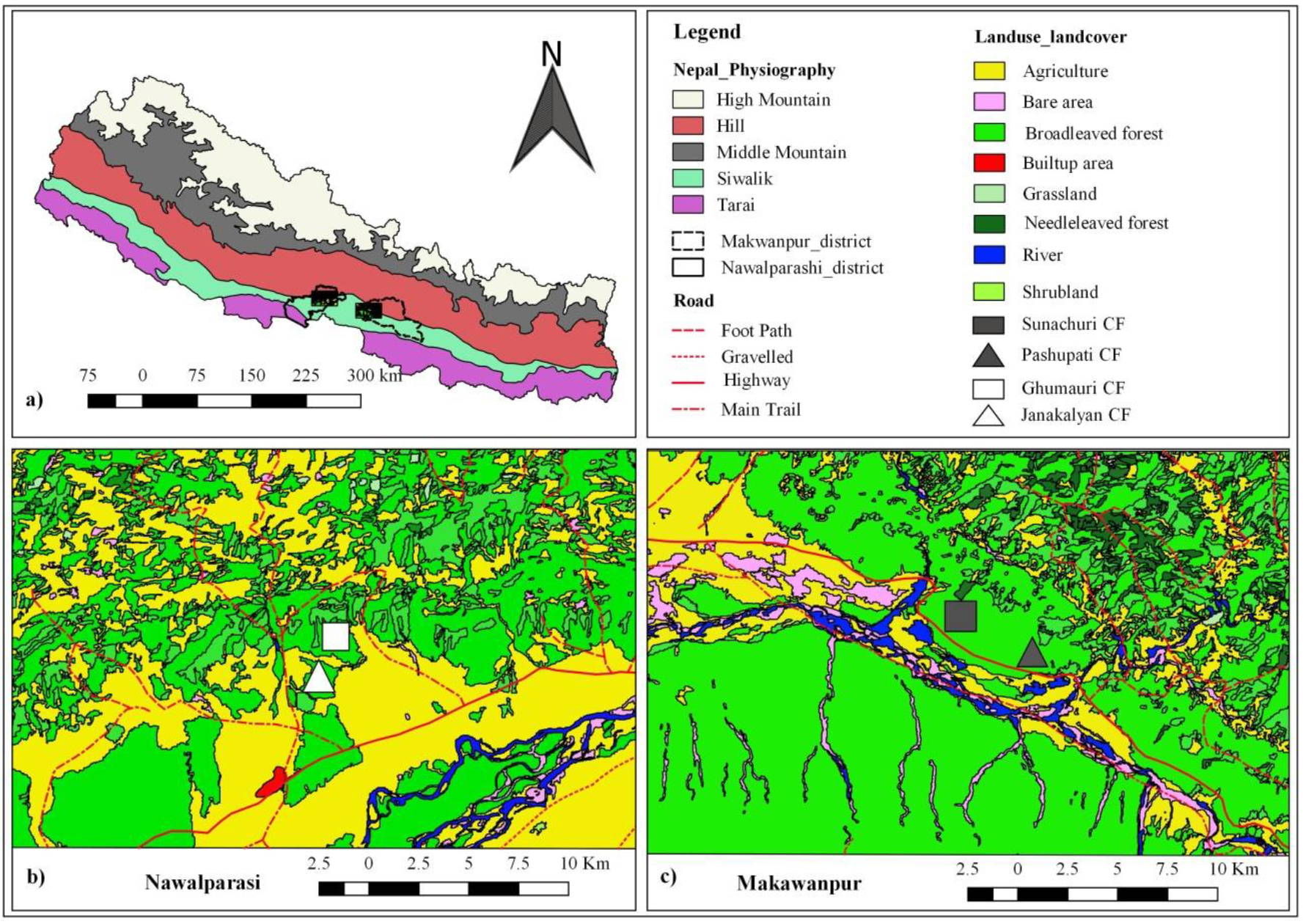
Maps showing the study areas; a) Location of Nawalparasi and Makawanpur district in the physiographic regions of Nepal, b) land cover of Nawalparasi site, c) land cover of Makawanpur site

All four of the forests in this study are located in the *Siwalik* (*Churiya*) hills. The Siwalik range is geologically young [give a date for this], and forms an east-west band of unconsolidated hills that runs parallel to the south of the Himalayas.

The sampled forests are located between 200-500 m altitude. The climate is subtropical and monsoonal, with hot and humid summers, and cool dry winters. Average annual rainfall is 2200 mm (1971–2010), of which 80% falls during the monsoon (June to August), and average annual temperature is 24.6 °C (2000-2010; CBS 2011). The forests in all the four sites is dominated by Sal (*Shorea robusta*). Sal is a member of Dipterocarpaceae, a tropical family mainly distributed in the Indo-Malayan region, and forms extensive mono-dominant or mixed forests in the southern part of the Himalayas, and in the tropical to subtropical climate of the Indian subcontinent (Gautam and Devoe 2006). Sal is a robust, gregarious, semi-deciduous tree species, and is an important high-value timber species extensively used in construction and furnishing. *Dillenia pentagyna* Roxb., *Buchanania latifolia* Roxb. and *Mallotus philippensis* (Lam.) Müll. Arg. are main sub-canopy species. *Clerodendrum viscosum* Vent. is the most common native shrub forming ground vegetation.

*Chromolaena odorata* (English name Siam weed and Nepalese name Seto banmara (‘White forest killer’), family-Asteraceae), is among the top 100 of the world’s worst invasive alien species (Lowe et al. 2000). Now it has spread in more than 100 countries in Asia, Europe and Africa and America and reported as problematic invasive weed in more than 35 countries (https://www.cabi.org/ISC/datasheetreport/23248). It is a light-demanding species and flourishes in disturbed forests, roadsides, fallow and abandoned lands. Its biological and morphological attributes (such as a long tap root and production of large quantities of wind dispersed seeds) give it a competitive advantage over native species (Joshi et al. 2006, Malahlela et al. 2015). *Chromolaena* can grow to three meter in height and forms a dense layer above the ground (Figure 2). This plant has already severely invaded the lowland districts of central and eastern Nepal at elevations below 1000 m and is now spreading into western districts (Tiwari 2005).

**Figure 2:**
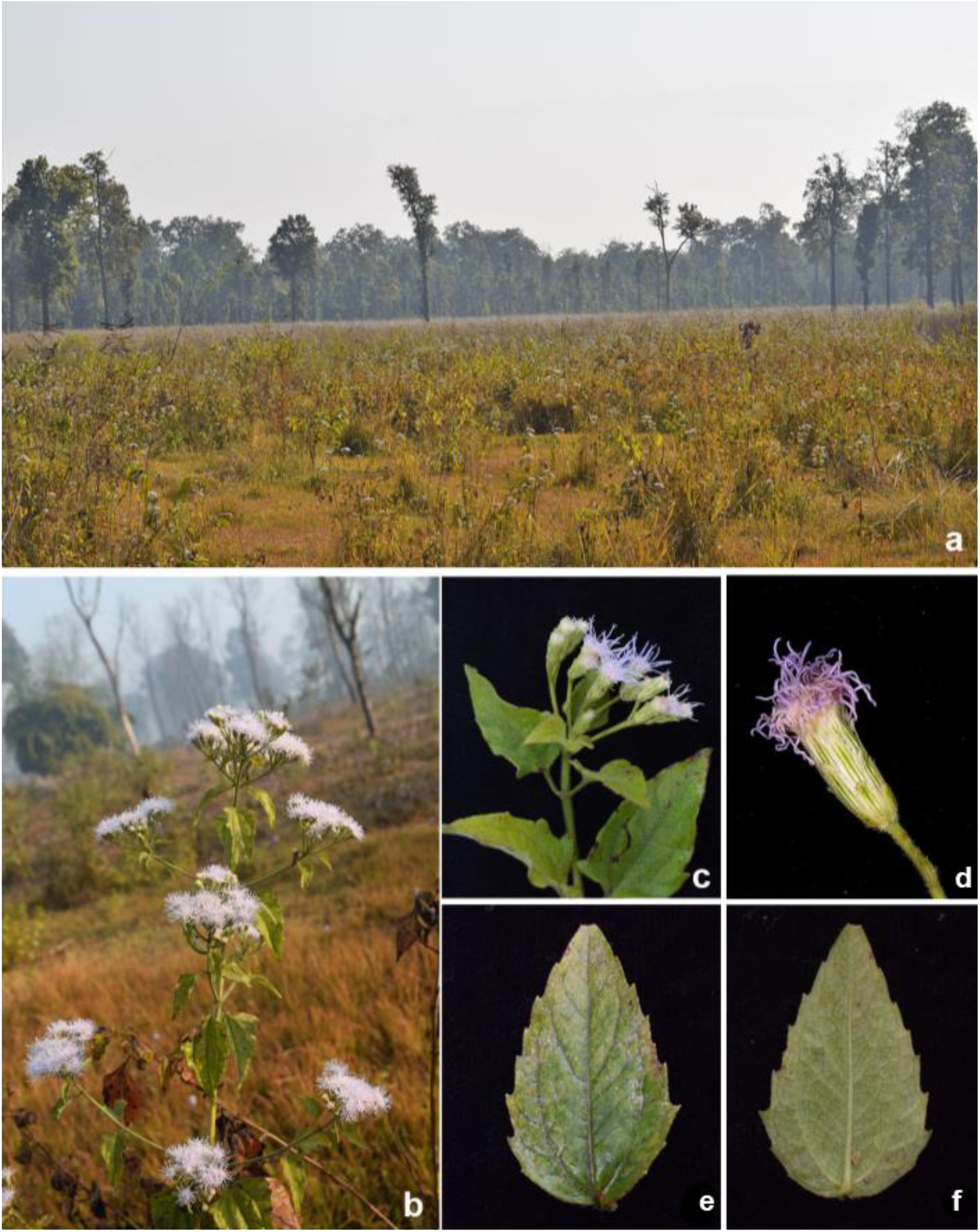
*Chromoleana* morphology, a-an open area invaded by the species; b-flowering branch; c-inflorescence detail; d-flower head detail; e & f-leaf dorsal and ventral surface showing margin and venation.

### Vegetation sampling

All the forests sampled were linked to a disturbance source, i.e. road or human settlement. We sampled vegetation along transects from the disturbance source into the forest interior. In each forest two transects were made. Before laying out the plot, the length of transect and number of plots were identified. The shortest distance between plots as between 100-150 m, depending on forest size, and in each forest 28 to 30 plots were sampled.

Vegetation data were collected in plots of 10 × 10 m. Each plot was divided into four subplots of 5 × 5 m. Diameter at Breast Height (DBH) of all the tree individuals greater than 5 cm DBH were measured in the plots. Canopy cover above the plot was measured using a spherical densitometer, with four readings were taken at each plot following the Lemmon (1956) protocol.

*Chromolaena* cover was estimated in all subplots by standing at the center of the subplot. The cover of subplots was combined to estimate cover for the 100 m^2^ plot. The same method was used to estimate shrub and herb cover.

In each plot two quadrats of 1 m^2^ were sampled randomly to record species richness. All herbaceous plants, shrubs and tree seedlings were recorded in each quadrat.

In each plot ground disturbance (grazing, tree/saplings lopping and trampling) was recorded on a scale of 0 to 3, where 0 represents absence of disturbance and 3 being severely disturbed. Distance of the plot from the nearest road was measured using Google Earth Pro.

To evaluate how CFUGs are managing *Chromolaena* in the study sites, we interviewed CFUG leaders (n=8, Chairman and Secretary in each CF) and one local knowledgeable person as indicated by the CF Chairman (n=4, one in each CF). Similarly we also interviewed CFUG leaders (Chairman or Secretary) in each of 15 other CFUGs in other parts of the country (Tanahu, Chitwan, Gorkha, Sindhuli and Jhapa district) which have Sal forest with *Chromolaena* invasion. Some informal discussions with local people were also conducted in public areas where people gather for each CF visited, to explore the general understanding of invasive species and their management.

### Data analysis

Ground disturbance was calculated combining three variables i.e. grazing, lopping and trampling, using Principal Component Analysis (PCA). PCA first axis score was used to represent ground disturbance complex. The predictor variables were checked for collinearity and only one of the collinear variables was selected for further analysis (see supplementary material). *Chromolaena* cover was our response variable. We used Zero Inflated Beta regression to evaluate impact of forest attributes on *Chromolaena* cover as our response variable is a proportion and contains many zeros (Bürkner 2017). Zero Inflated Beta regression is suitable when response variable is vegetation cover and consists of proportion data between zero and one (Keim et al. 2017). *Chromolaena* cover was modelled against each covariate individually and significant covariates were chosen. A full model analysis was run with *Chromolaena* cover as response and with all the non-collinear independent variables as predictors. Predictor variables that did not explain any variation in the model were subsequently dropped in the final model. Forest types were included as the random variable in the model. The R package BRMS (Bayesian Regression Model using ‘Stan’) (Bürkner 2017) was used for the regression analysis. The R^2^ for each model was calculated using *add_criterion* function of BRMS (Bürkner 2017). Each predictor variable was centered and scaled by subtracting its mean and dividing by its standard deviation prior to regression analysis so as to facilitate model convergence as well as to make relative effect size of predictor variables directly comparable (Muscarella et. al., unpublished). All analyses were performed in R version 3.5.3 (R Core Team 2019).

We compared the differences in *Chromolaena* cover among canopy cover classes using Analysis of Variance (ANOVA) and a post-hock Tuckey test. Canopy cover was categorized as low, medium and high. Values below the 1st quartile were considered low and those above 3rd quartile were considered high. Values lying around the median were considered as medium.

## Results

*Shorea robusta* was the most common species and the dominant canopy forming tree species in all the CFs studied. A total of 120 native plant species were recorded from those four forests. Native species richness ranged from one to 20 species per plot with a mean of 11.41 and standard deviation of 3.48. In addition to *Chromolaena odorata*, six other invasive species, namely *Spermacoce alata* Aubl., *Hyptis suaveolens* (L.) Poit., *Ageratum conyzoides* L., *Mimosa pudica* L., *Senna tora* (L.) Roxb. and *Mikania micrantha* Kunth. were also recorded. *Chromolaena odorata* was present in 60% of the plots with cover ranging from 0 to 95%.

### Correlation among stand attributes and *Chromolaena* cover

*Chromolaena* cover was negatively correlated with canopy cover, shrub cover, basal area and tree density. The strongest correlation was with forest canopy cover followed by basal area and tree density. Forest canopy cover in turn was positively correlated with basal areas and tree density. Shrub cover and native species richness were not correlated with *Chromolaena* cover. Native species richness was not correlated with any of the measured stand attributes (Table 1). Similarly, native species richness between invaded and non-invaded plots was not different (*t* = 0.64349, df = 112.34, *p*= 0.5212).

**Table 1:** Correlation matrix of forest stand attributes including Chromolaena cover and environmental variables

|  | Chrom<br>cover | Canopy<br>cover | Basal<br>area | Tree<br>density | Shrub<br>Cover | Native<br>richness | Herb<br>cover | Distance | Elev | Slope | Disturbance |
| --- | --- | --- | --- | --- | --- | --- | --- | --- | --- | --- | --- |
| Chrom. cover | 1 | -0.59 | -0.32 | -0.32 | -0.18 | 0.10 | 0.13 | -0.41 | -0.29 | -0.25 | 0.11 |
| Canopy cover | -0.59 | 1 | 0.48 | 0.58 | -0.00 | 0.04 | -0.03 | 0.26 | 0.23 | 0.06 | -0.13 |
| Basal area | -0.32 | 0.48 | 1 | 0.23 | 0.032 | -0.26 | 0.03 | 0.09 | -0.05 | -0.03 | -0.19 |
| Tree density | -0.32 | 0.58 | 0.23 | 1 | -0.01 | 0.03 | 0.01 | 0.11 | 0.09 | -0.01 | -0.20 |
| Shrub Cover | -0.19 | -0.00 | 0.032 | -0.01 | 1 | 0.00 | -0.10 | 0.03 | -0.12 | -0.08 | -0.07 |
| Native richness | -0.10 | 0.04 | -0.26 | 0.03 | 0.00 | 1 | -0.05 | 0.01 | 0.04 | -0.01 | -0.20 |
| Herb Cover | 0.13 | -0.03 | -0.03 | -0.01 | -0.10 | -0.05 | 1 | 0.01 | -0.08 |  | 0.04 |
| Distance | -0.41 | 0.26 | 0.09 | 0.11 | 0.03 | 0.08 | 0.01 | 1 | 0.54 | 0.01 | -0.22 |
| Elev | -0.29 | 0.23 | 0.03 | 0.09 | -0.12 | 0.30 | -0.08 | 0.54 | 1 | 0.49 | 0.02 |
| Slope | -0.25 | 0.06 | -0.05 | 0.04 | -0.01 | 0.19 | -0.01 | 0.21 | 0.49 | 1 | 0.00 |
| Disturbance | 0.11 | -0.13 | -0.19 | -0.20 | -0.07 | 0.16 | 0.04 | -0.22 | 0.02 | 0.00 | 1 |

### Environmental covariates affecting *Chromolaena* cover

Regression models containing canopy cover, distance to the road, shrub cover and slope had the highest mean r^2^ value, indicating these variables have impact on the abundance of the invasive species. *Chromolaena* cover declined linearly along the canopy cover gradient (Figure 3). Similarly *Chromolaena* cover linearly declined away from the road, with increasing shrub cover and slope (Supplementary figures 1, 2 and 3). Canopy cover had the largest effect size on *Chromolaena* cover, −0.53 (−0.85, −0.21) while it has relatively lower error for the regression estimates (Table 2). Distance to the road had the second largest effect on *Chromolaena* cover i. e. −0.35 (−0.64, −0.15). Slope and shrub cover had relatively smaller effects (table 2). Canopy cover, the most important stand attribute affecting *Chromolaena* cover, in turn increased with increasing distance from the nearest road (Figure 4).

**Figure 3:**
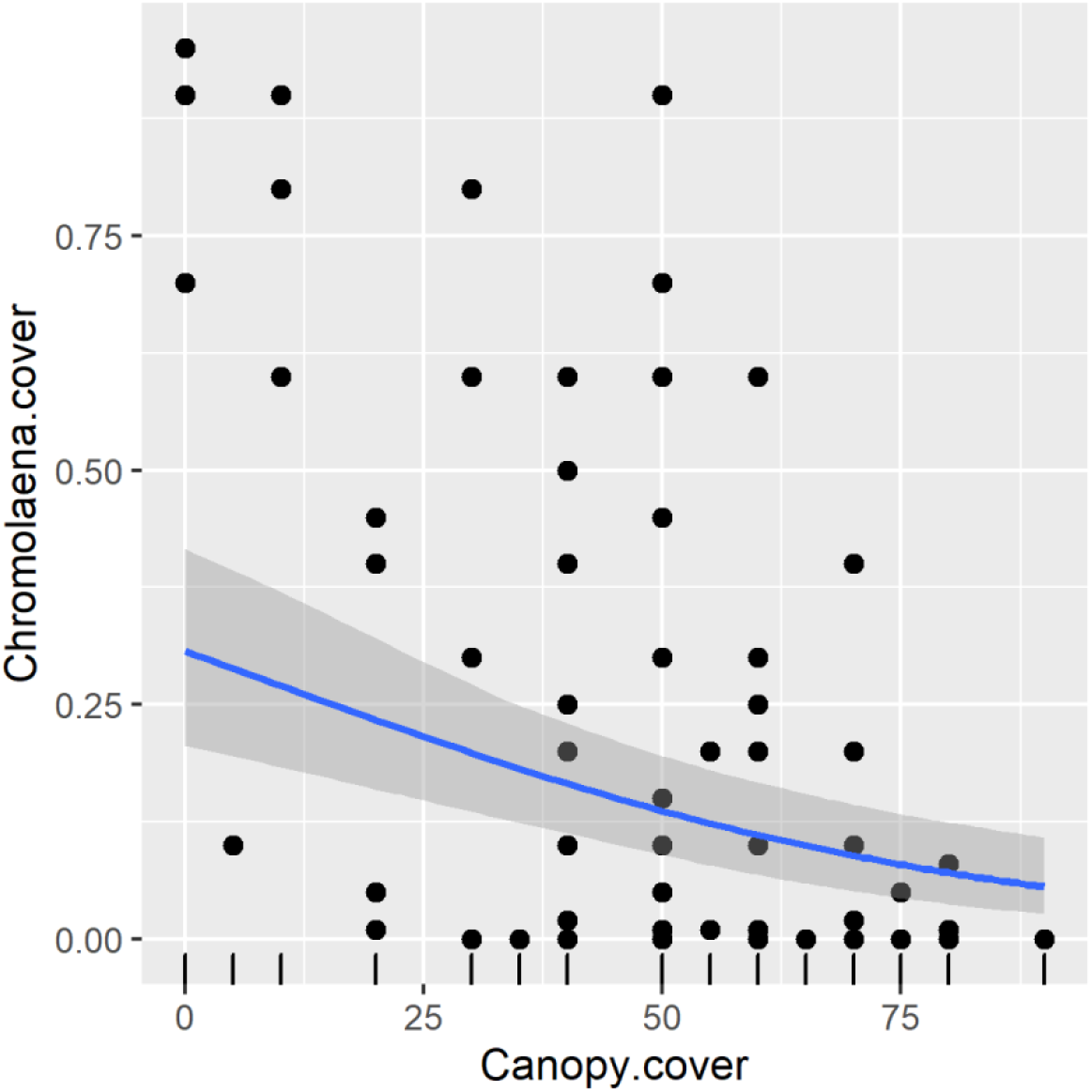
The relationship between *Chromolaena* cover and canopy cover showing fitted line and its 95% confidence intervals

**Figure 4:**
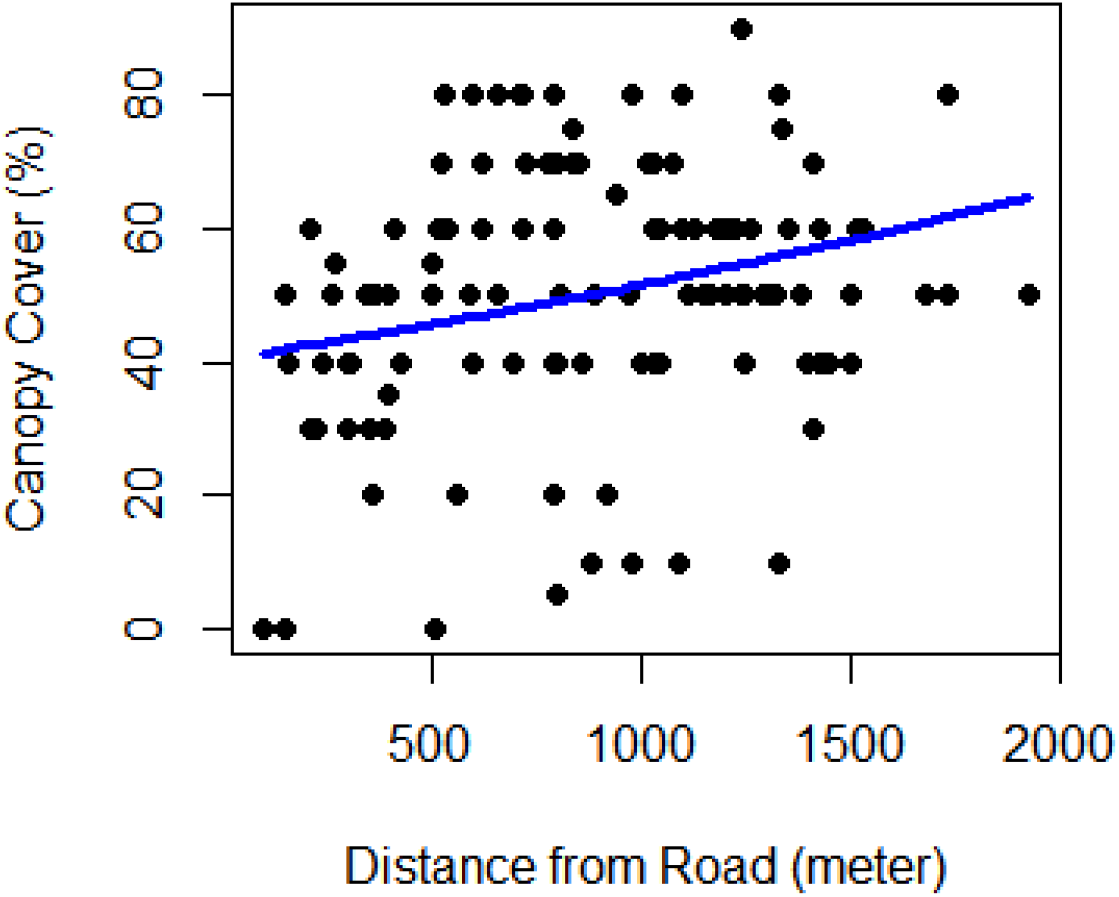
Relationship between canopy cover and distance to the nearest road.

**Table 2:** Model summary of Bayesian regression analysis where *Chromolaena* cover is response and other forest attributes are predictors.

| Variables | Estimate | Estimated error | 95% confidence interval |  |
| --- | --- | --- | --- | --- |
|  |  |  | Lower | Upper |
| Intercept | -1.15 | 0.23 | -1.64 | -0.68 |
| Canopy cover | -0.53 | 0.13 | -0.78 | -0.28 |
| Shrub cover | -0.21 | 0.14 | -0.47 | 0.06 |
| Slope | -0.28 | 0.13 | -0.54 | -0.01 |
| Distance | -0.29 | 0.14 | -0.56 | -0.02 |
| sd (Random effect of sites) | 0.29 | 0.32 | 0.01 | 1.15 |

*Chromolaena* cover did not show any trends with native species richness, herb cover and ground disturbance complex. The regression model indicates that the forest canopy cover was the overriding covariate affecting *Chromolaena* cover in sal forest (Ttable 2).

ANOVA, and post-hoc Tuckey test, showed that *Chromolaena* cover was different among the canopy cover classes. Cover was highest in forests with low canopy cover and lowest when canopy cover was higher (Figure 5, df=2, f=13.81, p<0.001). However, *Chromolaena* cover was not different between high and moderate canopy cover (Figure 4).

**Figure 5:**
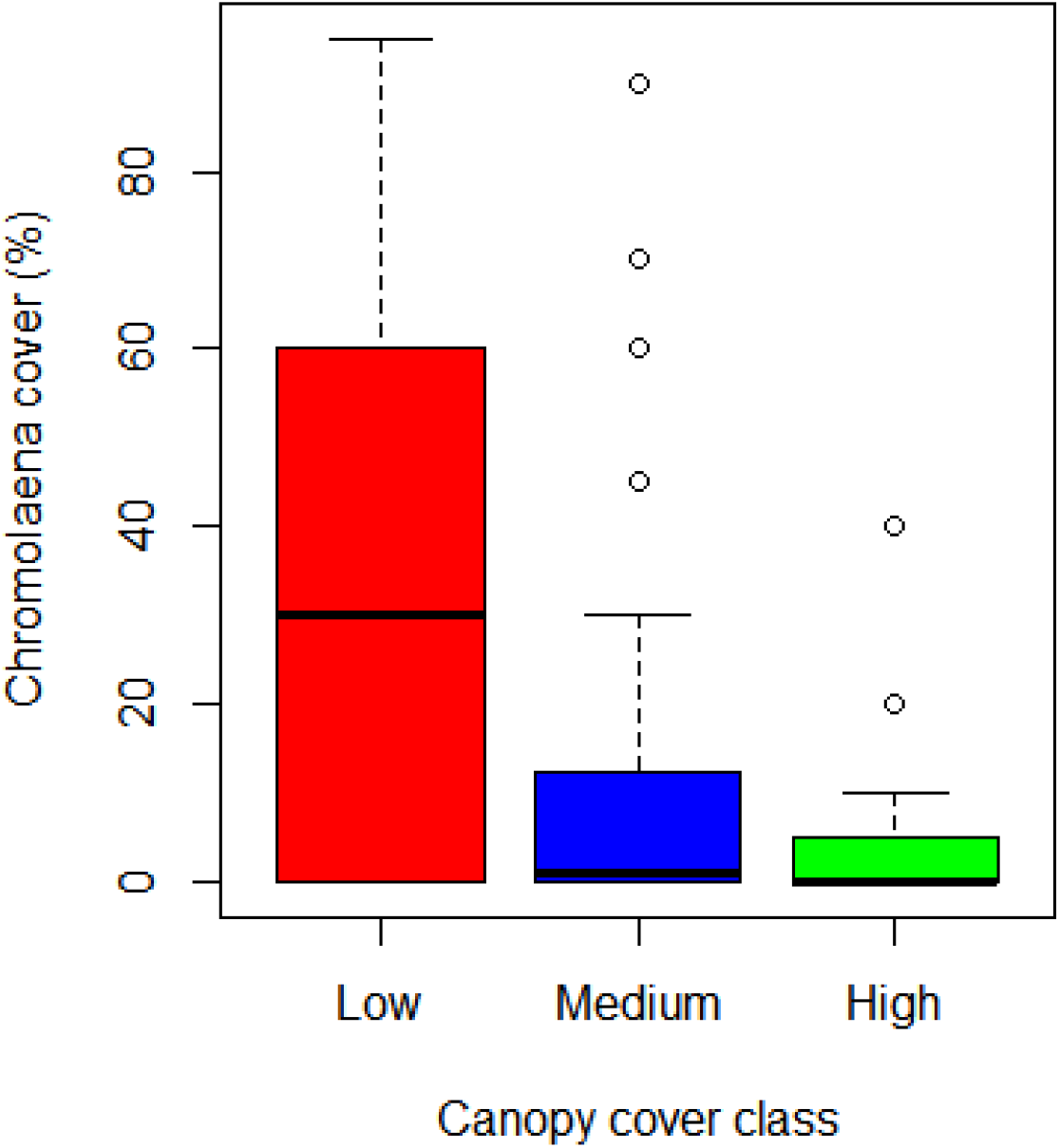
*Chromolaena odorata* cover in different canopy classes

We recorded that CFUGs are organizing regular bush clearing (*jhadi safai*) of understory plants in their CFs. However, these activities generally occur in forested areas and rarely in open parts invaded by *Chromolaena*. We found that CFUGs do not yet have specific programs targeting *Chromoleana* or other invasive plant; however, new management initiatives are being tested targeting these species.

## Discussion

The results demonstrate that forest stand attributes and other environmental variables affect the cover of *Chromoleana* in the Sal forests of Nepal. We discuss how attributes of Sal forests determine the cover of invasive species and highlight practical relevance of findings to managed community and other fragmented and disturbed forests.

### Canopy cover is the overriding covariate affecting *Chromolaena* cover

We have demonstrated that canopy cover, plot level disturbance, slope, and distance from the road all have some effect on *Chromolaena* cover in Sal forest. *Chromolaena* cover declines gradually away from the roadside which is probably due to high propagule pressure along roads. Roadsides in turn have lower canopy cover and more open areas and are important driver of invasion from local to the regional level (Flory and Clay 2006, Follak et al. 2018b). Roads bring propagules as well as create disturbances and open spaces (vacant niches), which consequently favour invasion. Roads provide corridors for invasive species, connecting them with suitable habitats, therefore, roadsides and forests edges often have high density of invasive species (Benedetti and Morelli 2017, Follak et al. 2018a).

Accessible parts of Sal forests are subjected to anthropogenic disturbances involving lopping trees, cutting saplings and trampling ground for firewood and fodder collection. Disturbance is an important variable affecting invasive species in forests. Counterintuitively, we found that ground disturbance had a very weak correlation with *Chromolaena* cover, and it did not improve the regression model, which indicates that ground disturbance is not a major factor governing the cover of *Chromolaena* in the forests in this study.

Canopy cover showed a negative relationship with *Chromolaena* cover in Sal forest. In the composite model containing disturbance, and distance to disturbance sources, forest stand level canopy cover is the overriding factor to control *Chromolaena* cover. *Chromoleana* cover declines linearly along increased canopy cover. In general, this negative relationship reiterates that *Chromolaena* is light demanding and prefers to grow in well illuminated areas (Joshi et al. 2006). In addition, as the invading species is an understory shrub it cannot compete with trees for light, consequently the canopy trees limit this crucial resource for *Chromolaena*. Joshi et al. (2006) also found that seed production of *Chromolaena* is suppressed in low light intensity. Higher canopy cover implies lower level of light availability below forest canopy. Many Invasive Alien Plant species (IAPS) prefer to grow in open areas in forests and forest ecotones (Mavimbela et al. 2018). Open areas in forests provide sites for regeneration and growth of IAPS and have higher proportion of IAPS density and coverage compared to closed-canopy areas (Charbonneau and Fahrig 2004, Driscoll et al. 2016). Nevertheless, the impact of canopy may also be dependent on the nature of invading species, as shade tolerant invasive species may be favoured where there is a dense canopy (Martin et al. 2009).

### Native species richness and *Chromolaena* cover

Conventional diversity resistance hypothesis asserts that sites with higher species richness have lower susceptibility to exotic invasions, mainly at local scale (Fridley et al. 2007). However, this hypothesis is not always supported by empirical studies (Peng et al. 2019); some studies corroborate (Kennedy et al. 2002) the hypothesis while others refute (Wiser et al. 1998). Alternatively, it is also argued that native species may even facilitates invasion (Fischer et al. 2009). Our study in the Sal forests shows that *Chromolaena* cover is not correlated with the higher levels of native biodiversity (species richness). Most of the published analysis of the effect of native richness on invasion comes from studies on grasslands (Kennedy et al. 2002), and diversity resistance experiments in forest systems are scarce. The main mechanism for invasion resistance is thought to be competition. It has been suggested that richness alone may not resists invasion rather there may be role of other factors co-varying with diversity which may contribute in invasion resistance of communities. In this case, forest canopy cover appears to be a more important factor than species richness with respect to community competitiveness to invasion resistance.

### Management implication

*Chromolaena* is one of the world’s worst invasive alien plant species (Lowe et al 2000). National policy documents categorize its impacts as ‘massive’ in Nepal (GoN, 2018), and its distribution in the Himalayas is expected to expand with climate change (Lamsal et al. 2018, Shrestha and Shrestha 2019). This species, along with other invasive species, demands immediate action so that their expansion to new location can be curtailed, the existing biomass can be controlled and ecological and biodiversity loss can be averted.

The Community Forestry program in Nepal is exemplary in restoring degraded forests and has played a key role in increasing forest cover and averting deforestation in Nepal (Niraula et al. 2013, Oldekop et al. 2019). Local people have also observed that *Chromolaena* abundance has been suppressed with forest restoration and canopy closure. Although CFs do not have specific plans and action to control *Chromolanena*, it appears that they have unwittingly played an important role in controlling *Chromolaena* in forests by protecting forest and increasing forest canopy. Control of invasive species through increased forest cover could be an ‘undocumented contribution’ of CFUGs of Nepal. However, additional data is needed from different physiographic regions and socio-economic settings to evaluate this hypothesis.

Our findings have immediate practical relevance in forest management. Nepal has more than 20,000 Community Forests and these forests are mostly small patches of forest interspersed with settlement and agriculture. These forest patches are subjected to disturbance associated with biomass extraction, grazing and forest silviculture and many of these CFs are potentially vulnerable to invasion by *Chromolaena*. CFs should consider enhancing forest canopy cover to suppress the growth of *Chromolaena*. Currently, Nepal has adopted intensive silviculture practices in Sal forests. Tree felled and canopy opened areas are highly susceptible to invasion by *Chromolaena* therefore such patches within forests should be monitored regularly to control invasion by *Chromolaena*.

The results of this study show that forest areas along the roadside have higher cover of *Chromoleana*. Intact forest margins along the roadside potentially buffer propagule dispersal towards forest interiors (Cadenasso and Pickett 2001). Therefore, increasing tree density and forest crown along road side could be a strategy to control the cover and control the spread of *Chromolaena* in fragmented forests. Forest managers should consider restoring degraded foress and increasing tree crown along roadsides and open areas so that invasive species can be suppressed while gaining other forest ecosystem services.

### Conclusion

Our study clearly indicated that forest canopy cover can resists invasion of *Chromolaena* in Sal forest. The resistance mechanism could be related to resource limitation, primarily light, to the invading species. Disturbance on the ground or undergrowth is probably not a primary driver facilitating invasion in forest when the invading species is light demanding. Our results provide practical insights for the management of Sal forests and degraded areas to avert invasion by invasive species, results that may apply to other forest types and other light demanding invasive species.

## Acknowledgements

We would like to thank Laxman Paudel, Chandra Bahadur KC, Ram Chandra and Raju Bhandari for their help during the field visits. This paper is one of the outputs of the Darwin Initiative UK funded project 23-031 ‘Science-based interventions reversing negative impacts of invasive plants of Nepal.

