## Supplemental file for "Forest canopy resists plant invasions: a case study of *Chromolaena odorata* in sub-tropical Sal (*Shorea robusta*) forests of Nepal"

**Supplementary figures**

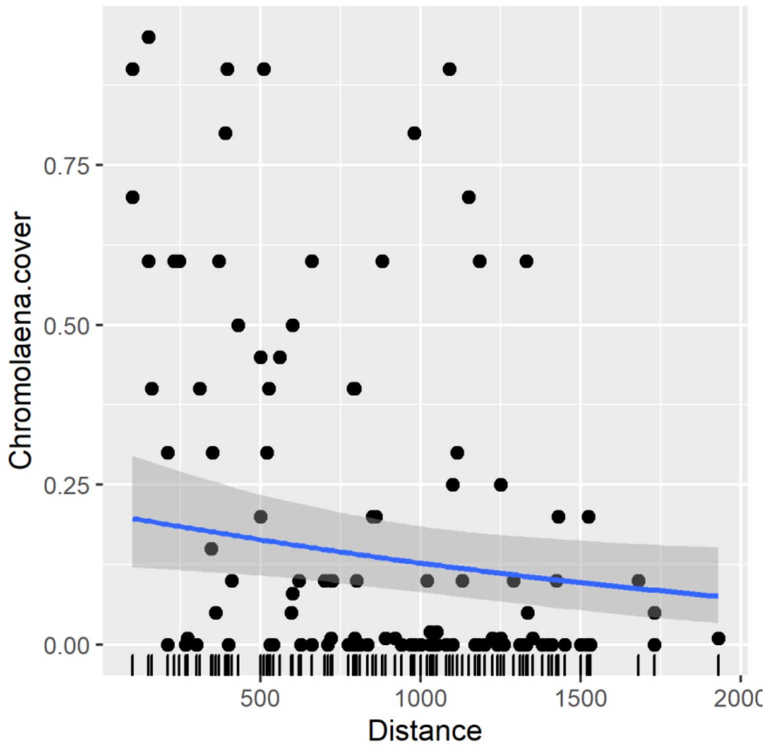


**Supplementary Figure 1**: The relationship between *Chromolaena* cover and distance from the nearest road showing fitted line and 95% confidence interval

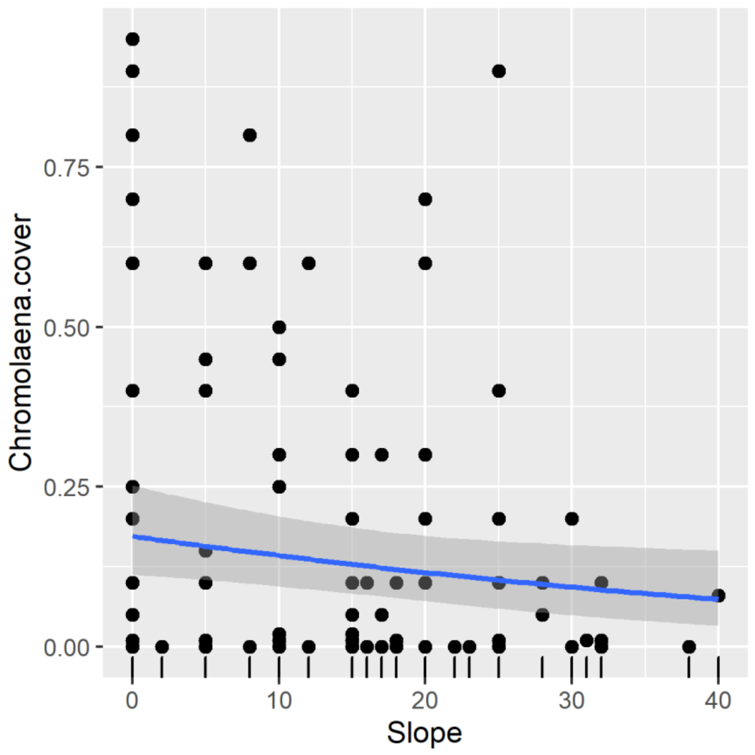


**Supplementary Figure 2**: The relationship between *Chromolaena* cover and slope showing fitted line and 95% confidence interval

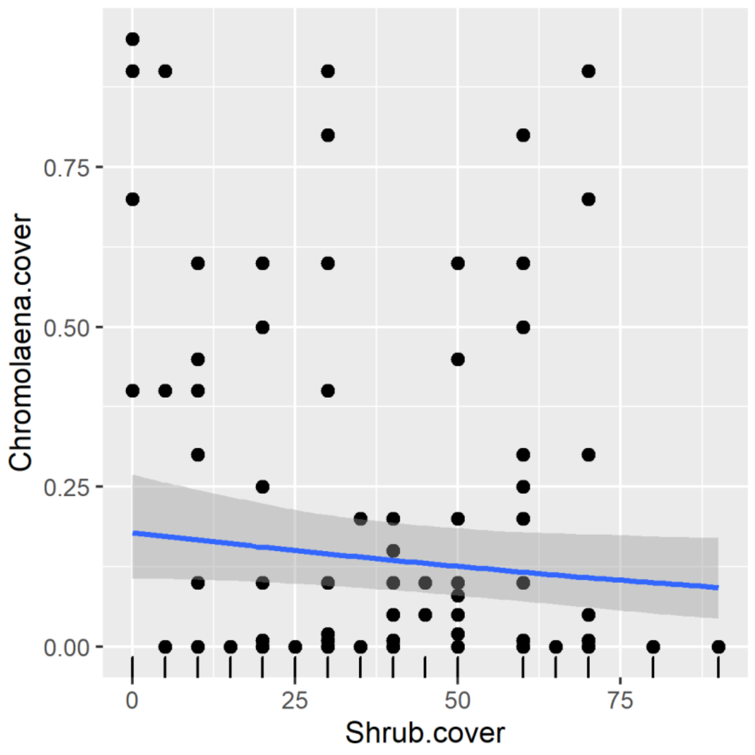


**Supplementary Figure 3**: The relationship between *Chromolaena* cover and shrub cover showing fitted line and 95% confidence interval
